# A *Bbs5* mouse model reveals pituitary cilia contributions to developmental abnormalities

**DOI:** 10.1101/2020.08.18.256537

**Authors:** Melissa R. Bentley, Staci E. Engle, Courtney J. Haycraft, Reagan S. Andersen, Mandy J. Croyle, Kelsey R. Clearman, Addison B. Rains, Nicolas F. Berbari, Bradley K. Yoder

**Affiliations:** Department of Cell, Developmental and Integrative Biology, University of Alabama at Birmingham, Birmingham, AL 35294; Department of Biology, Indiana University-Purdue University Indianapolis, Indianapolis Indiana, 46202

## Abstract

Primary cilia are critical sensory and signaling compartments present on most mammalian cell types. These specialized structures require a unique signaling protein composition relative to the rest of the cell to carry out their functions. Defects in ciliary structure and signaling result in a broad group of disorders collectively known as ciliopathies. One ciliopathy, Bardet-Biedl Syndrome (BBS; OMIM 209900), presents with diverse clinical features, many of which are attributed to defects in ciliary signaling during both embryonic development and postnatal life. For example, patients exhibit obesity, polydactyly, hypogonadism, developmental delay, and skeletal abnormalities along with sensory and cognitive deficits, but for many of these phenotypes it is uncertain which are developmental in origin. A subset of BBS proteins assembles into the BBSome complex, which is responsible for mediating transport of membrane proteins into and out of the cilium, establishing it as a sensory and signaling hub. Here we describe two new mouse models for BBS resulting from a congenital null and conditional allele of *Bbs5*. *Bbs5* null mice develop a complex phenotype including craniofacial defects, skeletal shortening, ventriculomegaly, infertility, and pituitary anomalies. Utilizing the conditional allele, we show that the male fertility defects, ventriculomegaly, and pituitary abnormalities are only found when *Bbs5* is mutated prior to P7 indicating a developmental origin. In contrast, mutation of *Bbs5* results in obesity independent of the age of *Bbs5* loss. Compared to other animal models of BBS, *Bbs5* mutant mice exhibit pathologies that suggest a specialized role for Bbs5 in ciliary function.

## Introduction

Primary cilia are microtubule-based structures that emanate from the surface of nearly every mammalian cell type. The ciliary membrane is enriched in a unique set of membrane proteins and signaling components that sets it apart from the cell membrane (1). This enrichment cultivates a highly specialized and responsive sensory and signaling hub for the cell. The accumulation of the proper signal transduction components at the ciliary membrane is crucial for cilia function and ultimately depends on the cooperation of several macromolecular machines, one of which is the BBSome. The BBSome is an octameric complex containing BBS1, BBS2, BBS4, BBS5, BBS7, BBS8, BBS9 and BBS18/ BBIP10 (2, 3). Interactions between Intraflagellar Transport Protein IFT22 (also known as RABL5) and BBS3 (also known as Arl6) are then responsible for the recruitment of the BBSome to the base of the cilium via interactions with the BBS1 subunit (4–6). The recruitment process is also aided by Rab8, the Rab8-specific GEF, Rabin8, and Rab11 (2, 3, 7). BBS5 is structurally and functionally unique based on predictions that it may directly mediate membrane interactions through its two plextrin homololgy (PH) domains capable of binding to phosphoinositides (2). Based on BBS5’s structure and physical interactions within the BBSome, it is unlikely that it is actually able to interact with the membrane via these PH domains (8). Thus, the functional role and importance of BBS5 in the BBSome remains poorly understood.

Bardet-Biedl Syndrome (BBS) patients exhibit a wide range of highly variable pathologies including but not limited to: obesity, hypogonadism, polydactyly, cognitive deficits, renal anomalies, and retinitis pigmentosa. To date, mutations in twenty-one different loci (BBS *1-21*) have been associated with BBS. Mutations specifically affecting the core BBSome complex represent a large proportion of BBS patients (9), with 2% of the mutations occurring in *BBS5* (10). Previously, congenital mutant mouse models of BBSome components BBS1, BBS2, BBS4, BBS7, and BBS8 have been described and recapitulate several, but not all, of the phenotypes associated with the clinical features of the disorder. Additionally, a conditional allele for *Bbs1* has been described with phenotypes that recapitulate some of the clinical features (11–15). However, work done thus far in *Bbs5* models has been limited and only demonstrated minor retinal degeneration (16, 17). We sought to assess the pathophysiology of *Bbs5* loss of function alleles using congenital and conditional *Bbs5* mutant approaches. Our goal was to distinguish between phenotypes that are developmental in origin from those that occur as a consequence of loss of BBS5 functions needed for tissue homeostasis in adults. To accomplish this goal, we analyzed phenotypic consequences of Bbs5 disruption during development, in juvenile, and adult stages. We report phenotypes including: submendelian survival ratios, shortened skeletons, craniofacial defects, sterility, obesity, ventriculomegaly, persistence of the buccohypophyseal canal, and pituitary gland abnormalities. Out of these, obesity was unique in that it is the only phenotype seen in both the congenital allele and when *Bbs5* loss is induced after postnatal day 7, suggesting roles for *Bbs5* in both development and adult that can impact energy homeostasis. The phenotypes observed in *Bbs5* mutant mice described here are directly related to the pathologies presented by BBS patients, and provide the first whole animal validation of the *Bbs5* mutant mouse model as a valuable tool to further understand the molecular mechanisms resulting in the pathologies common to BBS.

## Materials and Methods

### Generation of Bbs5 mutant alleles

All animal studies were conducted in compliance with the National Institutes of Health *Guide for the Care and Use of Laboratory Animals* and approved by the Institutional Animal Care and Use Committee at the University of Alabama at Birmingham. Mice were maintained on LabDiet^®^ JL Rat and Mouse/Irr 10F 5LG5 chow. *Bbs5* knockout first (Bbs5^tm1a(EUCOMM)Wtsi/+^; *Bbs5^-/+^*) embryonic stem cells, from C57BL/6NTac background mice, were obtained from Eucomm and injected into C57BL/6J (JAX Stock No: 000058) blastocysts to establish the *Bbs5^-/-^* (*tm1a*) line. The allele was then maintained on the C57BL/6J strain. *Tm1c* conditional allele mice were generated by mating *tm1a* to FlpO recombinase mice (C57BL/6J) thus removing the LacZ and Neo cassettes and generating a conditional allele (*tm1c; flox*). Progeny that contained the recombined allele were crossed off of the FlpO line and bred to Cagg-Cre^ERT2^ males (C57BL/6J) to generate the *tm1d* (delta) allele. Here we refer to these alleles as the *tm1a* (*Bbs5^-/-^*), *tm1c* (*Bbs5^flox/flox^*) and *tm1d* (*Bbs5^Δ/Δ^*) alleles. (**Figure 1A**). Primers used for genotyping are as follows for the *tm1a* allele: 5’-TTCAGTTGGTCAGTTTTGTATCGT-3’, 5’-TCAGCACCGGATAACAGAGC-3’, and 5’-CATAGTTGGCAGTGTTTGGGG-3’ and for the *tm1c* and *tm1d* alleles:5’-TGTTTTGTTGGTAGATGATGCATGGG-3’, 5’ CAGAGAAGCATTGGTAATAACCGAGC-3’, 5’-TGAGGGTAGGAACGGAGCTCAGAG-3’.

**Figure 1.**
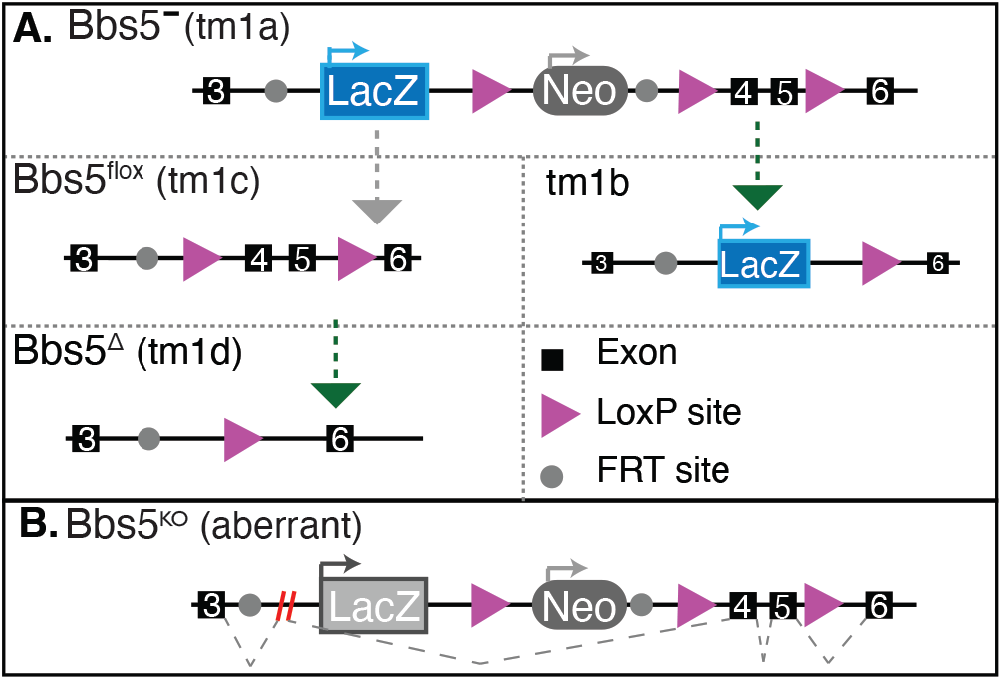
Mouse Alleles. A) The knockout allele construct depicting the congenital knock out allele (*tm1a*), floxed allele (*tm1c*), and recombined alleles (*tm1b* and *tm1d*). Exons are depicted as black boxes, *LoxP* sites as purple arrows and *FRT* sites as grey circles. Grey arrows indicate that a FlpO mouse was used to generate the subsequent allele. Green arrows indicate that a Cre-expressing mouse was used to generate the subsequent allele. B) The *tm1a* allele, as verified by sequencing of the resulting cDNA, results in an alternatively spliced allele excluding both the LacZ and Neo coding regions.

### Embryo Isolation

Timed pregnancies using *Bbs5^+/-^* animals were established with embryonic time-point of E0.5 being noted at noon on the morning of observing the copulatory plug. To isolate embryos, pregnant females were anesthetized using isoflurane followed by cervical dislocation. Embryonic tissues or whole embryos were isolated and fixed in 4% paraformaldehyde (Sigma PFA, 158127) in PBS.

### Mouse embryonic fibroblast (MEF) Isolation

Embryos were isolated at E13.5. Following the removal of the liver and head, embryos were mechanically dissociated and cultured in DMEM (Gibco, 11039-021) supplemented with 10% Fetal Bovine Serum, 1X Penicillin and Streptomycin, 0.05% Primocin, 3.6μl/0.5L β-mercaptoethanol. Cells were grown to confluency at which time media was changed to DMEM containing 0.5% FBS to induce cilia formation.

### Tissue Isolation and Histology

Mice were anesthetized with 0.1 ml/ 10 g of body weight dose of 2.0% tribromoethanol (Sigma Aldrich, St. Louis, MO) and transcardially perfused with PBS followed by 4% paraformaldehyde (PFA; Affymetrix Inc., Cleveland, OH). Tissues were post-fixed in 4% PFA overnight at 4°C and then cryoprotected by submersion in 30% sucrose in PBS for 16–24 hours then cryosectioned for immunofluorescence and Hematoxylin (Fisher Chemical, SH26-500D) and Eosin (Sigma-Aldrich, HT110132-1L) staining was performed.

### Immunofluorescence microscopy

Ten (10) μm tissue sections (brain sections were 35 μm) were used for immunofluorescence microscopy. For staining MEFs, cells were grown on glass cover slips treated with 0.1% gelatin until confluent, then serum starved using DMEM containing 0.5% FBS for 24 hours to induce cilia formation (18). Sections were fixed with 4% PFA for 10 minutes, permeabilized with 0.1% Triton X-100 in PBS for 8 minutes and then blocked in a PBS solution containing 1% BSA, 0.3% TritonX-100, 2% (vol/vol) normal donkey serum and 0.02% sodium azide for one hour at room temperature. Primary antibody incubation was performed in blocking solution overnight at 4°C. Primary antibodies include: Acetylated a-tubulin (Sigma, T7451) direct conjugated to Alexa 647 (Invitrogen, A20186) and used at 1:1000, ACIII (Encor, CPCA-ACIII, 1:1000), Arl13b (Proteintech, 1771-1AP, 1:500), CCSP1 (Abcam, ab40873, 1:250), Mchr1 (Invitrogen, 711649, 1:1000), PECAM1 (Abcam, ab7388, 1:250), and SPC1 (Millipore Corp, AB3786, 1:250). Cryosections were then washed with PBS three times for five minutes at room temperature. Secondary antibodies diluted in blocking solution were added for one hour at room temperature. Secondary antibodies included: Donkey conjugated Alexa Fluor 647, 488, and 594 (Invitrogen, 1:1000). Samples were then washed in PBS and stained with Hoechst nuclear stain 33258 (Sigma-Aldrich) for 5 minutes at room temperature. Cover slips were mounted using SlowFade Diamond Antifade Mountant (Life Technologies) for PVN and ARC sections and Immu-Mount (Thermo Scientific) for all others. Brain sections were imaged on a Leica SP8 confocal using 60X objective (NA=1.4). All other fluorescence images were captured on Nikon Spinning-disk confocal microscope with Yokogawa X1 disk, using Hamamatsu flash4 sCMOS camera. 60x apo-TIRF (NA=1.49) or 20x Plan Flour Multi-immersion (NA=0.8) objectives were used. Images were processed using Nikon’s Elements or Fiji software.

### Skeletal Preparations and bone measurements

The skin and internal organs (except brain) of 2-month-old mice were removed and the skeletons were submerged in 1% KOH overnight at room temperature. Skeletons were rinsed and cleaned of further excess tissue and fresh KOH solution added. Skeletons were left in KOH solution until sufficient tissue could be removed. Skeletons were rinsed with water and placed in a solution of 1.6% KOH and 0.004% alizarin red for two days. Skeletons were rinsed with water and placed in clearing solution (2 volumes glycerol; 2 volumes 70% ethanol; 1 volume benzyl alcohol). Skeletons were then stored in 100% glycerol and imaged using a Nikon SMZ800 stereo microscope.

### Tamoxifen Cre Induction

Recombination of the *tm1c* allele was induced in juvenile *Bbs5^flox/flox^*; *CAGG-cre^ERT2^* mice at postnatal day 7 by a single intraperitoneal (IP) injection of 9 mg tamoxifen (Millipore Sigma, T5648) per 40 g body weight. Tamoxifen was dissolved in corn oil. Adult animals were induced at 8 weeks old by IP injections of 6 mg/40 g (body weight) tamoxifen, administered once daily for three consecutive days.

### Sequencing

Fluorescence based Sanger sequencing using the Illumina NextSeq500 Next Generation Sequencing (NGS) instrument at the Heflin Center for Genomic Sciences was performed on cDNA generated from brain, heart, lung, kidney, testes, and retinal extract in wild-type and *Bbs5^-/-^* mice.

### MRI imaging

Magnetic Resonance Imaging (9.4T) of post-mortem brains was conducted using T2 weighting (TE: 36 TR:1800). Imaging was performed on adult mice at two months of age. All imaging was performed at the UAB Small Animal Imaging Shared Facility. Images were analyzed using Horos and ImageJ software.

### Statistical Analysis

Calculations were performed using Graphpad Prism and Microsoft Excel. Specific tests used are indicated in figure legends with significance indicated as follows: * p≤0.05, ** p≤0.01, *** p≤0.001

## Results

### Bbs5^-/-^ mice have decreased viability but no defects in ciliogenesis

We first sought to verify that *Bbs5* is widely expressed using the *LacZ* cassette engineered into the *Bbs5^-/-^* allele (**Figure 1A**). However, we were unable to detect β– galactosidase staining in any tissue. We then investigated the expression of the targeted allele by RT-PCR in several tissues. Using primers located upstream of the cassette and within the *LacZ* gene, we were unable to detect a product. We therefore checked for expression of the *Bbs5* transcript using primers located in exons upstream and downstream of the cassette. Surprisingly, we identified a single transcript produced. Subsequent sequencing showed that the transcript produced from the *tm1a* allele uses an alternative splice site within the engineered exon and joins with exon four of the *Bbs5* gene, splicing around the *LacZ* coding sequence, explaining the lack of β–galactosidase staining. The aberrant transcript contains several in frame translational termination codons early in the sequence and is therefore predicted to be a null, or severe hypomorphic allele (**Figure 1B**).

Homozygous mutant mice (**Figure 1A;** *Bbs5^-/-^*) are viable but exhibit a significantly increased mortality by weaning age (P21) compared to heterozygous and wild type littermates (**Figure S1A**). Our studies indicate that during the final stages of embryonic development, E18.5-birth, all genotypes are present at the ratios expected from heterozygous matings (*χ*^2^(2, N=47; 7 litters)=3.09, p> 0.05 progeny (**Figure S1A**)). However, Mendelian ratios reflect a significant reduction (*χ*^2^(2,N=141, 23 litters)=19.93, p< 0.001) in the observed number of mutant animals at weaning, (**Figure S1A**) indicating failure to thrive and perinatal lethality. By immunostaining for Arl13b and acetylated α-tubulin, there were no overt differences detected in number or length of primary cilia in analyzed tissues, indicating that *Bbs5^-/-^* mutant mice do not display a general defect in ciliogenesis. *Bbs5^-/-^* cells form cilia at a similar frequency and with similar lengths as controls (**Figure S1B, C and D**).

BBS patients can present with highly variable phenotypes. This is thought to be related to the modifying effects of individual patients’ different genetic backgrounds. In mice, it has been reported that phenotypes associated with mutations in other *Bbs* genes are also affected by genetic background (19). Previous reports of background-dependent lethality in BBS mutant mice have been attributed to neural tube closure defects and pulmonary developmental defects (20, 21). During embryo isolations, we never observed neural tube closure defects and Mendelian ratios were observed after neural tube closure (E18.5-birth). For these reasons, we went on to assess whether pulmonary developmental defects could be contributing to perinatal lethality in *Bbs5^-/-^* mice. Histological analysis of lungs at E18.5 do not show obvious differences in alveolar space or pulmonary interstitium (**Figure S2A**). Immunofluorescence staining for the alveolar type I cells, vasculature, and alveolar type II cells using antibodies against SPC1, PECAM1, and CCSP1, also did not reveal differences compared to control littermate lungs (**Figure S2B, C** and **D**). Thus, in contrast to other BBS mutant models, perinatal lethality in *Bbs5^-/-^* mice is not associated with overt pulmonary defects.

Observationally, perinatal *Bbs5^-/-^* mice appear smaller. Similar to what has been observed in other BBS mouse models, growth retardation occurs during the first three weeks in mutant animals, allowing them to be easily distinguished from their littermates; this is possibly caused by the inability to nurse due to anosmia (15).

### Fertility defects in Bbs5 mutant animals

To determine if *Bbs5^-/-^* mutant mice were fertile, we performed homozygous by heterozygous matings. While both male and female heterozygous mice are fertile, no litters were produced when either the male or female was homozygous for the *Bbs5* mutant allele, indicating that both male and female *Bbs5^-/-^* mice are infertile. In other mouse models of BBS, infertility was associated with a lack of flagellated sperm (12). To investigate whether this could be the cause of the infertility in male *Bbs5^-/-^* mice, we isolated the testes and performed histological staining. In *Bbs5^-/-^* testes, no flagellated sperm were visible (**Figure 2A**). Furthermore, extraction of sperm from the epididymis of mutant mice also did not yield flagellated sperm, while isolation from wild-type and heterozygous animals did (**Supplemental video 1**). This could be a result of defects in flagella formation, sperm differentiation, or puberty defects related to disruption of the hypothalamic-pituitary-gonadal axis (22, 23).

**Figure 2.**
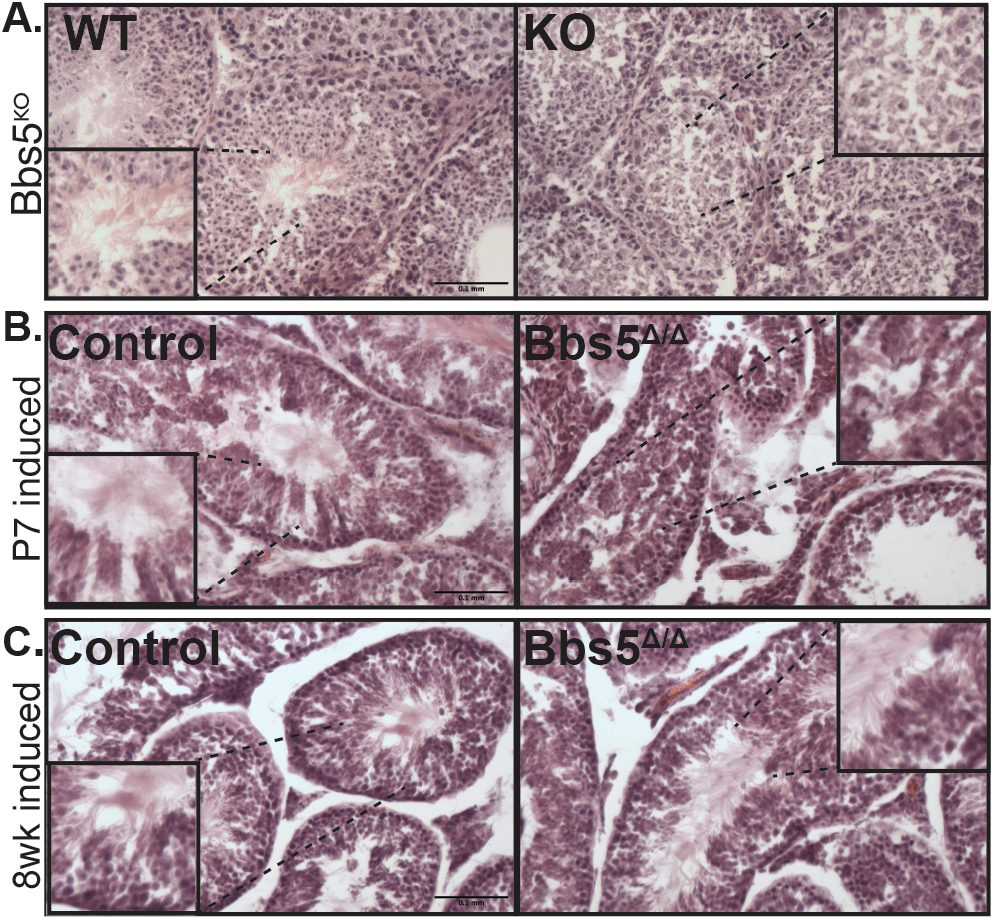
Testes Analysis. H&E staining of testes in A) Wild-type and *Bbs5^-/-^* mice, B) Juvenile induced conditionals and C) adult induced conditionals. Staining shows the presence of flagellated sperm in WT, *Bbs5^f/f^*, and adult induced *Bbs5^Δ/Δ^* animals versus the lack of flagellated sperm in *Bbs5^-/-^*and juvenile induced *Bbs5^Δ/Δ^* mice. (Scale bar 0.1mm)

To further evalaute whether this is a defect in develoment of the sperm versus maintanence, we utilized conditional *Bbs5^flox/flox^*; Cagg-Cre^ERT2^ animals and induced *Bbs5* loss at either 7 days (P7) prior to sexual maturation or after at 8 weeks of age. Testes isolated from *Bbs5^Δ/Δ^* mice that had been induced at P7 and analyzed at least two months post induction showed a variable phenotype, where 5 out of 8 male mice did not develop flagelated sperm and 3 mice did develop flagelated sperm. In 2 out of 3 of these mice, the number of flagelated sperm appeared reduced (**Figure 2B**). In the adult-induced (8 weeks) mutants analyzed 10 weeks post induction, flagellated motile sperm were present in all *Bbs5^Δ/Δ^* mice analyzed (N=5) (**Figure 2C**). These data indicate a developmental role for BBS5 during early spermatogenesis events, but not in sperm flagella maintenance.

### Bbs5 Mutant Obesity and Neuronal Cilia

As indicated, approximately one week following birth, surviving *Bbs5^-/-^* animals can be distinguished from their littermates due to their smaller size (data not shown). Over time, the surviving *Bbs5^-/-^* mutants not only catch up to their littermates with regards to body weight, but surpass them and become obese. To determine if the obesity observed in *Bbs5^-/-^* mutants was due to a developmental phenotype or a role for Bbs5 in adult homeostasis, we utilized the *Bbs5* conditional allele (*Bbs5^flox/flox^*). Using the near ubiquitously expressed Cagg-Cre^ERT2^ allele which has produced obesity phenotypes in other ciliopathy alleles, (24, 25) we analyzed adult phenotypes upon the conditional loss of BBS5 at P7 and 8 weeks of age. Both male and female conditional mutant *Bbs5^Δ/Δ^* animals become obese on breeder chow diet (10% crude fat) compared to their Cre negative controls (**Figure 3A)**.

**Figure 3.**
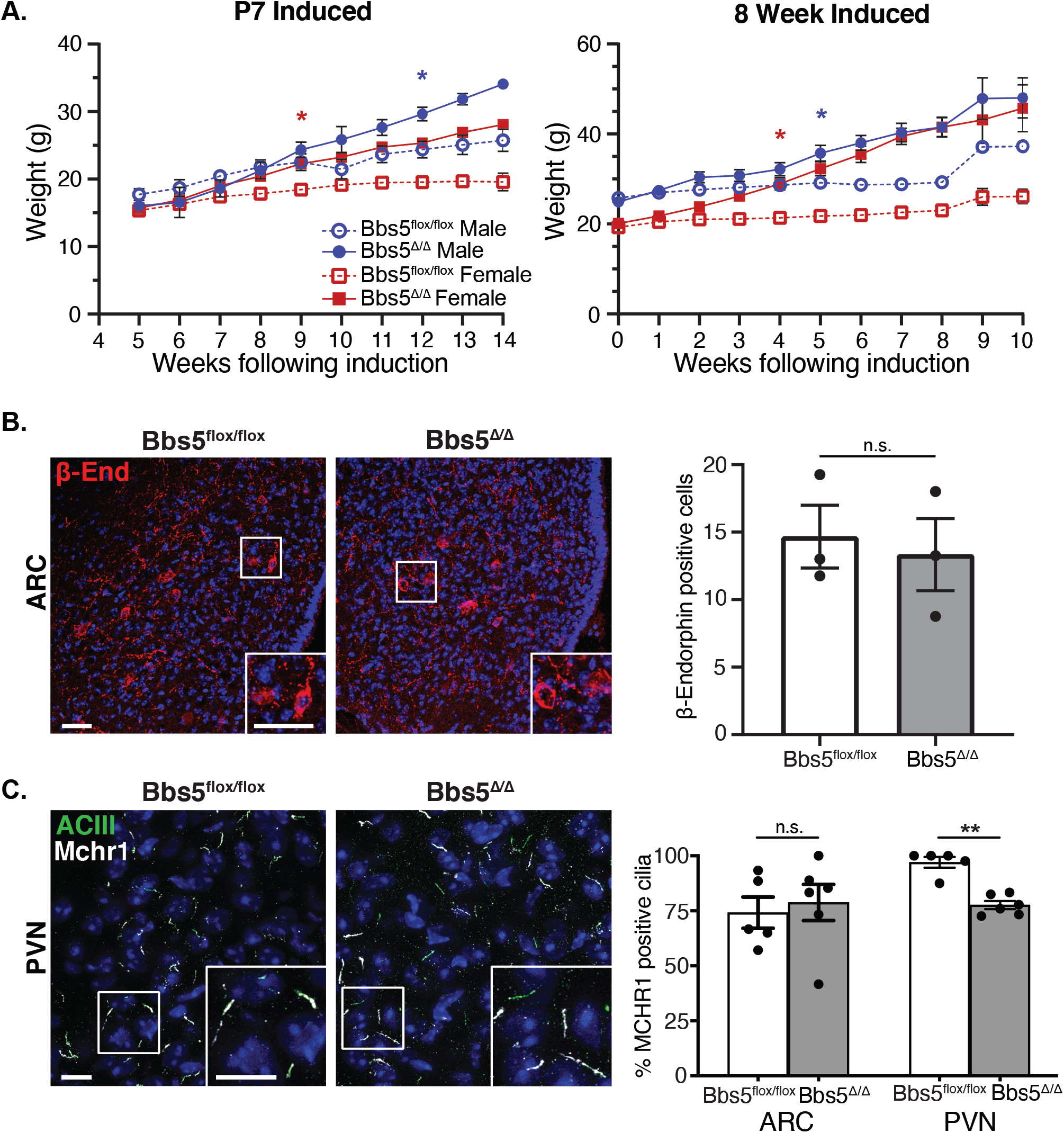
Obesity and Neuronal Cilia. A) Body weight measurement after conditional loss of Bbs5. (Left graph) Weights of male and female *Bbs5^f/f^* and *Bbs5^Δ/Δ^* mice following induction on postnatal day 7, N = 4 ♂ & 3♀ controls and 3♂ & 2♀ mutants. (Right graph) weights following adult induction at 8 weeks old (Adult Induced), N = 9 ♂ & 7♀ controls and 8♂ & 12♀ mutants. Asterix represent initial significant differences (P<0.05) using a mixed-effects analysis with multiple comparisons. Error bars represent SEM B) POMC neuron immunofluorescence in the Arcuate Nucleus (ARC) for β-endorphin (β-end, red) in control and adult induced mutant males (*Bbs5^Δ/Δ^*). (Right graph) Number of β-endorphin positive cells per section of ARC was not significantly different (n.s.) between genotypes in three males per group using a Students T-test. Scale bar 10μm. N = 3 control and mutant ♂ C) Primary cilia immunofluorescence for cilia marker adenylate cyclase III (ACIII, green) and cilia GPCR, melanin concentrating hormone receptor 1 (Mchr1, gray) in the Paraventricular Nucleus (PVN). (Right graph) Quantification of ACIII and Mchr1 double positive cilia in ARC and PVN revealed no significant differences in the ARC (n.s.) but reduced double positive cilia in PVN were observed using Student T-test (P < 0.01). Scale bar= 10μm. N = 3 ♂ & 2 ♀ controls and 3 ♂ & 3 ♀ mutants. All Hoechst stained nuclei blue. * p≤0.05, ** p≤0.01, *** p≤0.001

Other congenital BBS mutant mouse models develop obesity and display loss of POMC neuron labeling within the arcuate nucleus of the hypothalamus. This is consistant with either a loss of POMC neurons or a defect in leptin responsiveness in these mutants (26). However, in *Bbs5^Δ/Δ^* mutant mice, immunoflourescence for the POMC neuronal marker β-endorphin did not reveal differences between controls and *Bbs5^Δ/Δ^* mutants in cell numbers (**Figure 3B**) suggesting that the POMC neruonal population is intact and that there are no changes in cell number following to the onset of obesity. This is similar to what has been observed in other conditional cilia models and BBS mutants suggesting the loss of POMC neurons is due to alterations during neuronal development (27).

Both BBS2 and BBS4 are important for proper ciliary localization of G-protein coupled receptors like Melanin Concentrating Hormone Receptor 1 (MCHR1), which plays a role in feeding behavior and metabolism (28). Surprisingly, unlike *Bbs2* and *Bbs4* congenital knockout mice, *Bbs5^Δ/Δ^* mice still localize MCHR1 to the cilium in the hypothalamus (**Figure 3C**)(28). While in the arcuate nucleus (ARC), MCHR1 is found in cilia at comparable levels to controls, Mchr1:ACIII double positive cilia are significantly reduced in the paraventricular nucleus (PVN) of *Bbs5^Δ/Δ^* mice compared to controls (p=0.0024) (**Figure 3C**). We did not observe overt differences in the frequency or length of the cilium in the *Bbs5^Δ/Δ^* compared to controls. Together, these data suggest that changes in ciliary composition and subsequent signaling may initiate the obesity phenotype in adults and is not solely due to developmental patterning of the hypohthalmaus in this ciliopathy model.

### Decreased endochondral bone length in Bbs5 mutant mice

Surviving *Bbs5^-/-^* congenital mice displayed skeletal abnormalities as measured by an overall decrease in length from the tip of the nasal bone to the pubic symphysis (**Figure 4A**). This difference is present in both *Bbs5^-/-^* (p<.001) and *Bbs5^-/+^* (p<.001) mice compared to wild-type littermates. A decrease in the length of long bones, as represented by shortened femurs (**Figure 4B**) follows a similar trend in both *Bbs5^-/-^* (p<0.001) and *Bbs5^-/+^* (p<0.001) mice compared to wild-type littermates. During our gross inspection of the skeletons, we also noted that none of the *Bbs5^-/-^* mutant mice analyzed exhibited polydactyly. While this phenotype is commonly observed in human BBS patients, it has not been observed in any BBS mutant mouse models to date (11–13).

**Figure 4.**
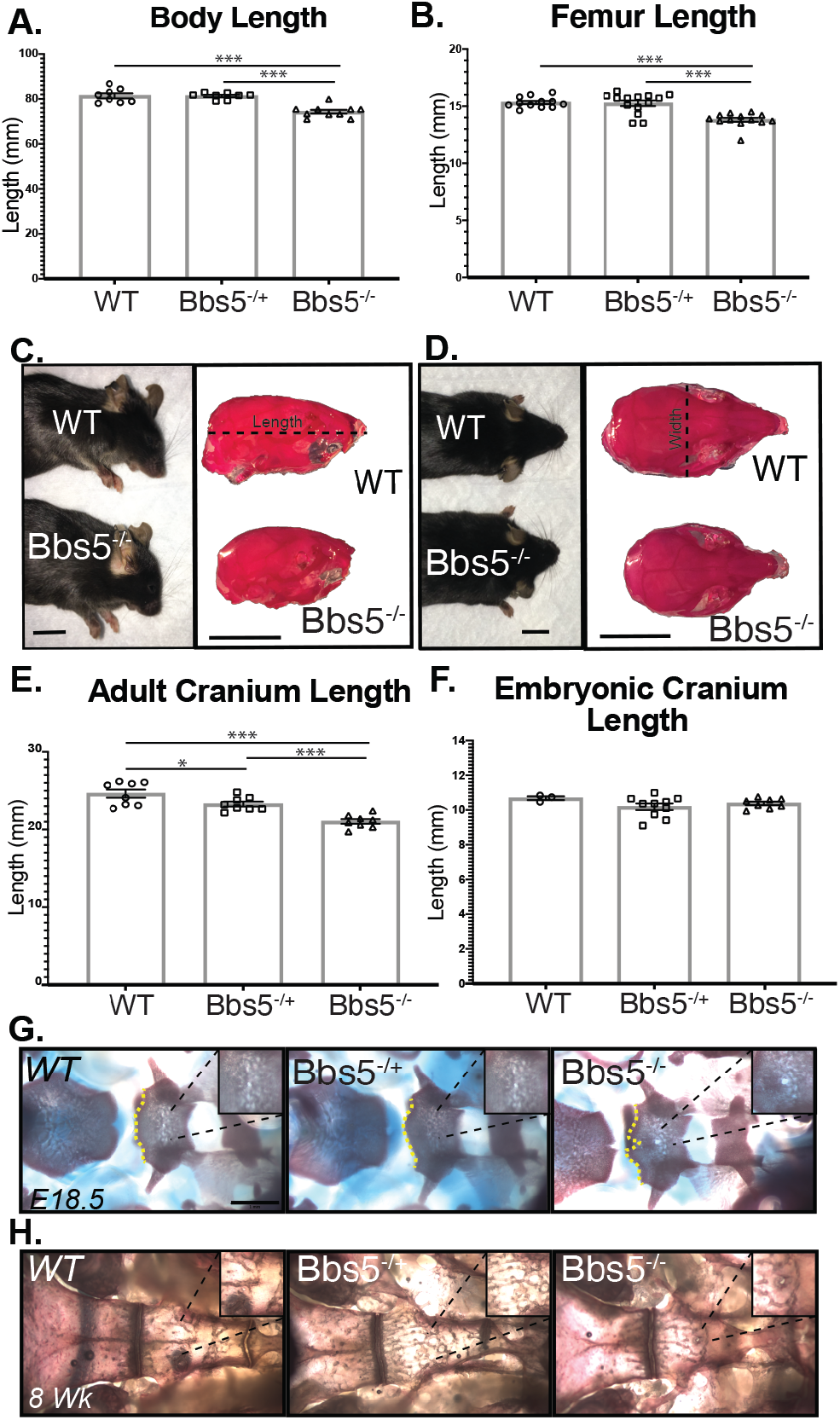
Skeletal Analysis. *Bbs5^-/-^* mice exhibit craniofacial and skeletal abnormalities. Measurements of and A) skeleton length N = 8 controls, 8 heterozygotes, and 10 mutants. and B) femur length, N= 12 control, 14 heterozygous, and 13 mutant femurs at 8 weeks old. C) Sideview of WT (top, left) and *Bbs5^-/-^ (*bottom, left) animals and skulls of WT (top, right) and *Bbs5^-/-^ (*bottom, right) that have been stained with alizarin red. (scale bar= 1mm). D) Overhead view of WT (top, left) and *Bbs5^-/-^ (*bottom, left) animals and skulls of WT (top, right) and *Bbs5^-/-^ (*bottom, right) that have been stained with alizarin red. (scale bar =1mm). E) Cranium Lengths in 2 month old WT, *Bbs5^-/+^*, and *Bbs5^-/-^* animals, N=8, 8, and 8 respectively. F) Cranium Lengths in E18.5 WT, *Bbs5^-/+^*, and *Bbs5^-/-^* animals, N= 3,10, and 8 respectively. Alizarin red and alcian blue staining of WT, *Bbs5^-/+^*, and *Bbs5^-/-^* cranial base (dorsal aspect) at G) E18.5 and H) 2 months old. For measurements, length was measured from the back of the scull to the tip of the nasal bone (dotted line in panel c). Error bars represent Standard error. Significance was determined via unpaired Ttest. * p≤0.05, ** p≤0.01, *** p≤0.001

### Bbs5^-/-^ animals postnatally develop shortened craniofacial bones

Craniofacial abnormalities in adult *Bbs5^-/-^* mice were observed in the skull (lateral view **Figure 4C**, overhead view **Figure 4D**). Skull length in adult mice measured from the tip of the nasal bone to the back of the skull is significantly different among the genotypes. For example, between wild-type *Bbs5^+/+^* and knockout *Bbs5^-/-^* animals, the distance is shorter in knockouts (p<0.001). Interestingly, we also observe significant differences between wild-type (*Bbs5^+/+^*) and heterozygous (*Bbs5^-/+^*) animals (p<0.05) and between *Bbs5^-/+^* and *Bbs5^-/-^* (p<0.001) (**Figure 4E**). These phenotypes were not present in E18.5 skulls analyzed, suggesting a role for Bbs5 in later stages of cranial development and growth (**Figure 4F**). These data are similar to previous reports of craniofacial abnormalities in other BBSome mutant animals. (13, 29)

Skeletal analysis also revealed structural abnormalities and a persistence of the buccohypophyseal canal in the basisphenoid bone at the base of the skull in E18.5 *Bbs5^-/-^* embryos (**Figure 4G**) and in adult *Bbs5^-/-^* animals (**Figure 4H**). These phenotypes are not observed in *Bbs5^-/+^* or wild-type mice. The buccohypophyseal canal is an ancestral vertebrate structure that typically disappears in mammals during development to generate a barrier between the pituitary gland and the oral cavity. The persistence of this canal was also described in *Gas1* knockout animals, which show reduced Sonic Hedgehog (Hh) signaling at the midline, along with pituitary development abnormalities in *Ift88* conditional (Wnt-1Cre), *Ofd1*, and *Kif3a* cilia mutant mice. This was attributed to altered regulation of the Hh signaling pathway in the midline of the associated embryos (30). Until now the only other reported case of basisphenoid abnormalities in BBS mice has been in BBS3/Arl6 congenital mutant models, which is not a member of the core BBSome complex (19).

### Brain and pituitary abnormalities in Bbs5 mutant mice

A potential cause for the smaller size of the *Bbs5^-/-^* mutant mice, along with defects in sperm production and abnormal bone length could be pituitary gland dysfunction. (31) This possibility is supported by the persistence of the buccohypophyseal canal in *Bbs5^-/-^* mice as well as BBS patients presenting with pituitary abnormalities (32). To further assess the pituitary gland in the *Bbs5^-/-^*, we performed magnetic resonance imaging (MRI) on heads of control and congenital mutants as well as conditional mutants where Bbs5 loss was induced early (P7) and in adults (8 week old) (**Supplemental video 2**). Sagital cross-sections of MRI images indicate that the mutant pituitary glands exhibit abnormal morphology with ectopic expansions caudally which were never observed in wild-type control littermates (3/5 mutant animals, **Figure 5A**). Importantly, histological analysis of sections through the pituitary gland in mutants that did not show structural abnormalities by MRI, revealed cellular abnormalities such as irregular boundaries and hyperplastic expansion between the Pars Intermedia (PI) and Pars Distalis (PD) regions. These abnormalities would not be identifiable by MRI analysis (**Figure 5B** and **5C**) and include irregular boundaries between the PI and Pars Distalis (PD) with neoplastic growths exhibited in the PI. Immunofluorescence staining using an antibody to the small GTPase Arl13b indicates that the wild-type Pars Nervosa (PN) region is sparsely ciliated, or lack Arl13b positive cilia, but the PI and PD are heavily ciliated (**Figure 5D**). In *Bbs5^-/-^* mutants the PI shows a reduction in Arl13b staining compared to wild-type (**Figure 5D**). The PD region *Bbs5^-/-^* mutant MRI analysis indicates that the pituitary glands in *Bbs5^-/-^* mutant mice are also significantly smaller (**Figure 5E**, p≤0.01). Future studies of these pituatary abnormalities may reveal how they contribute to the clinical features of BBS.

**Figure 5.**
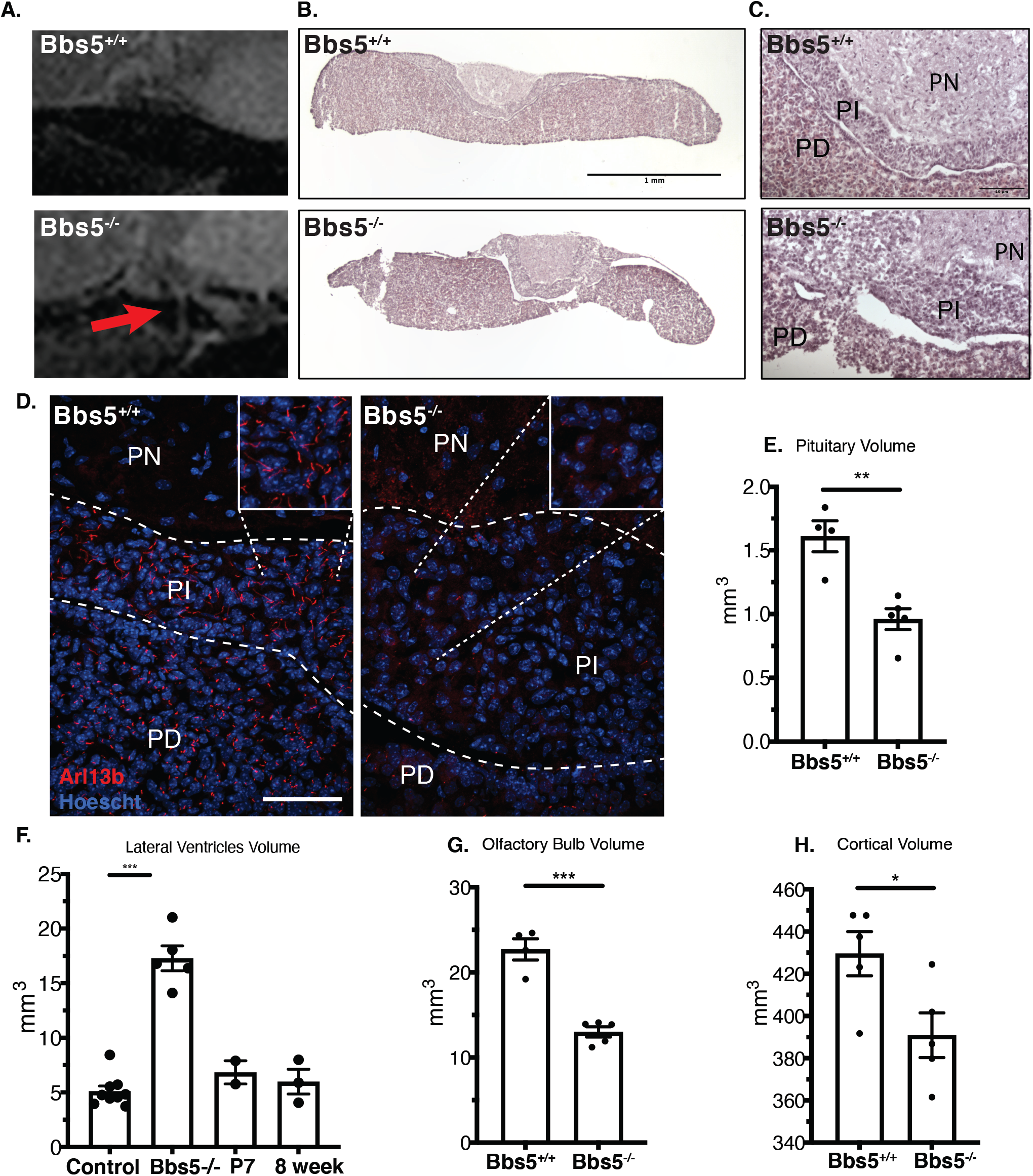
A) Sagittal cross section of pituitary MR images reveals structural abnormalities in *Bbs5^-/-^* animals compared to controls (red arrows). B) H&E histology of the pituitary (scale bar = 1mm), C) magnified H&E staining of the PN, PI and PD regions of the pituitary (scale bar =10 μm). D) Immunofluorescence staining of cilia in the pituitary using the small GTPase ArlI3b (scale bar =50 μm). PN= Pars Nervosa, PI= Pars Intermedia, and PD= Pars Distalis. Volumetric analysis of MR Images shows a significant change in: E) Pituitary, F) lateral ventricles of KO mice compared to control and juvenile and adult induced animals: (Control includes 5 wild-type and 4 *Bbs5^f/f^* animals, G) the olfactory bulb, H) cortex. Significance was determined via unpaired Ttest. * p≤0.05, ** p≤0.01, *** p≤0.001

Similar to other BBS models (12), adult (2-4 months old) *Bbs5^-/-^* mice exhibit the characteristic ventriculomegaly with an increase in the volume of the lateral ventricles (p<0.001). Interestingly, conditional mice that have been induced at either juvenile or adult timepoints, and imaged 2 and 4 months following Cre induction respectively, ventriculomegaly was not observed. (**Figure 5F**). In *Bbs5^-/-^* mice, the overall volume of the olfactory bulb (**Figure 5G**, p<0.001) and cortex (**Figure 5H** p<0.05) are reduced. These data suggest that some of the neural antaomical phenotypes observed in BBS are due to their roles in early postnatal development and not in adult homeostasis.

## Discussion

Classic BBS-associated obesity is observed in these animals. Most BBS obesity studies have been performed in congenital models. However, this study specifically utilizes a conditional allele for neuronal receptor localization studies. This allows for the interpretation of the consequences of BBS5 loss independent of developmental defects. Due to the fact that obesity occurs when induced at both juvenile and adult time points, it can be determined that obesity is driven by a process that occurs throughout the lifespan of the animal. Furthermore, the observations that POMC neuron number remains normal, cilia number is unaffected, and MCHR1 is trafficking appear normal (except minor defects in the PVN) in the hypothalamus distinguishes the conditional model from other BBS congenital mutant models (26, 28). These data suggest that obesity is being driven by alternative mechanisms than what have been proposed previously. Follow up studies will focus on comparing the congenital BBS5 mutant obesity with the conditional BBS5 mutant obesity to determine if loss of BBS drives the obesity phenotype through the same mechanism.

Congenital loss of BBS5 consistently results in a lack of flagellated sperm. Similarly, loss of BBS5 in a juvenile mouse shows a mixed impact on sperm flagellation. In contrast, disruption of BBS5 in adults has no impact on spermatogenesis. Based on these data, the fertility defects observed in male mutant mice are likely to be the consequence of developmental abnormalities. Spermatogenesis is dependent on several mechanisms including, but not limited to, proper neuronal signaling to coordinate Gonadatropin Releasing Hormone (GnRH) release followed by Folicle Stimulating hormone (FSH) and the proper function of the hypothalmus-pituitary-gonadal axis for proper tissue autonomous regulation of FSH and androgens (33). Cilia have been shown to play a role in regulating neuronal activity of GnRH neurons. The cilia on these neurons express the Kisspeptin receptor (Kiss1r), which is responsible for responding to kisspeptin and regulating the onset of puberty (22). The resulting animals that form flagellated sperm may be a result of the timing and efficiency of induction during a critical window in the initial wave of spermatogenesis. The high turnover rate of sperm production would indicate that, if BBS5 is necessary for spermatogenesis, its loss should affect sperm formation regardless of age of induction. Instead we note that loss of BBS5 in adult animals does not affect sperm production. This supports a role for BBS5 during initial spermatogenesis but not directly in flagella formation. These observations regarding fertility, paired with the skeletal abnormalities at the cranial base, and pituitary abnormalities point to hormonal dysregulation as a potential culprit driving the phenotypes observed in *Bbs5* mutant mice.

In addition to being smaller in size, pituitary glands in three out of five of these mice have defects that are visible by MRI analysis. The remaining two have defects visible following histological analysis. This result points to the possibility that pituitary dysfunction in these animals may be a result of defects in the developmental process itself. The observation that the primary cilia in *Bbs5^-/-^* pituitaries are also affected indicates that there may be further hinderance of pituitary function that is a direct result of ciliary signaling dysfunction, although this awaits more detailed analysis.

Overall our studies highlight several requirements for BBS5 in regulating the development of the axial and craniofacial skeleton. While craniofacial abnormalities have been reported in mouse models of BBS (13, 29), the only other reported case of basisphenoid abnormalities, as we observe in *Bbs5* mutants, is in *Bbs3/Arl6* congenital mutant models (19). This canal is hypothesized to be reminiscent of the transient developmental structure, Rathke’s pouch. During mammalian pituitary development the basal diencephalon gives rise to neuroectoderm, which along with oral epithelium, migrates via Rathke’s Pouch through the developing palatine bone to form the anterior pituitary. In contrast, the posterior pituitary, is derived from the neural ectoderm (34). In addition to the observation that the pituitary in *Bbs5^-/-^* mutant mice are structurally compromised compared to wild-type animals, points to possible hormonal dysregulation in these mutant mice. Defects in pituitary hormonal regulation could also underlie the developmental defects such as bone length and reproductive abnormalities observed in *Bbs5^-/-^* mutant mice. Furthermore, pituitary abnormalities such as hypoplasia, small Rathke’s cleft cyst, and pituitary enlargement have recently been reported in the BBS patient population (35). Thus, the BBS5 model described here will be a good model in which to explore the hypothalmus-pituitary-gonadal axis defects associated with disruption of the BBSome.

The development of the pituitary is an event that requires the tightly regulated synchronization of interactions between and migration of both the neural ectoderm and Rathke’s pouch derived from the oral ectoderm. It has been shown that abnormalities in the development of the pituitary can result in the persistence of the buccohypophyseal canal (30). In work done by the Dupé lab, there is a similar persistence of the buccohypophyseal canal in mice that are haploinsufficient for *Sonic Hedgehog (Shh*) (36). This becomes more severe in animals that are heterozygous for both *Shh* and the Notch pathway gene, *Rbpj*. These data indicate a requirement for both Shh and Notch signaling in closing of the buccohypophyseal canal. This points to the developing pituitary as a unique region within the embryo that is sensitive to the level of activity of the Shh and Notch pathways combined. It is widely accepted that canonical Hh signaling is dependent on the presence of the primary cilium. *Bbs5* mutant animals do not exhibit classic Hh signaling defects (e.g. dorsal ventral neural tube patterning defects, polydactyly) suggesting that it is largely unaffected in most of the embryo. This study suggests that the loss of Bbs5 specifically in the developing pituitary may be just enough to predispose animals to subtle Hh-associated pituitary abnormalities. This is further supported by disruption of Arl13b signaling in the intermediate region of mutant pituitaries, as Arl13b is also known to regulate Shh signaling events (37, 38). Of course, this result does not indicate whether cilia are still present, but unable to traffic Arl13b, or that cilia are absent altogether from the Pars intermedia in mutant mice. Attempts to answer this question included using traditional ciliary markers for ACIII, IFT components, and Acetylated α-tubulin were unsuccessful due to lack of expression of ACIII in the pituitary and difficulty getting the remaining antibodies to work in neuronal tissues. Based on the current understanding of BBSome function, it would be unlikely that the cilium is not present. Alternatively, a loss of cilia in the Pars intermedia could be a result of cell differentiation abnormalities, which may cause variability in cell types that may or may not be ciliated normally. Further investigation into the role of the primary cilium and the BBSome in pituitary development is necessary to definitively answer these questions.

By performing MRIs on congenital and conditional *Bbs5* mutant mice, we were not only able to identify structural abnormalities in the pituitary, but also to further expand on the classic BBS phenotype, ventriculomegaly. Based on the MRI data, the *Bbs5^-/-^* mice also have a reduction in cortical and olfactory bulb volume. MRIs performed on both juvenile and adult induced conditional *Bbs5* mutant animals addressed whether these phenotypes are a result of developmental consequence of loss of BBS5 or a requirement for BBS5 in normal tissue function. Conditional ablation of *Bbs5* at both juvenile and adult stages does not appear to result in enlarged ventricles.

In summary, the Bbs5 mutant mouse described here will be a good model to evaluate multiple phenotypes associated with BBS patients. Importantly, this includes pituitary defects. Pituitary abnormalities have been reported in both BBS and Joubert Syndrome (JBTS; OMIM 213300) patients (35, 39). This study is the first to show defects in pituitary development in a BBS mouse model. While not yet considered one of the classic pathologies associated with BBS or other ciliopathies, perhaps some of the underlying pathologies in patients are driven by a dysfunctional pituitary. Indeed, pituitary abnormalities have been noted in a study of a small number of BBS patients (35). This could also explain why mutation of *Bbs5* results in tissue specific phenotypes, which is unexpected given that *Bbs5* is thought to be expressed in all ciliated cells. Differences between this this model and other mouse models of BBS may provide evidence that mutations to BBS5 specifically target the pituitary. This evidence provides valuable insight into the mechanisms driving the disease state and may provide critical opportunities for pituitary-focused clinical intervention.

## Abbreviations

MEF: Mouse Embryonic Fibroblasts
ARC: Arcuate Nucleus
BBS: Bardet-Biedl Syndrome
IFT: Intraflagellar Transport
SHH: Sonic Hedgehog
ACIII: Adenylate cyclase III
MCHR1: Melanin Concentrating Hormone Receptor 1
PVN: Paraventricular Nucleus
PN: Pars Nervosa
PI: Pars Intermedia
PD: Pars Distalis
MRI: Magnetic Resonance Imaging

## Acknowledgements

We would like to thank the members of Dr. Bradley K. Yoder’s and Dr. Nicolas F. Berbari’s, laboratories for intellectual and technical support on the project. We would like to thank Dr. John Totenhagen, Dr. Anna Sorace and the support of the Small Animal Imaging Shared Facility (UAB). We would like to thank Dr. Sally Camper and her laboratory in the Department of Human Genetics at The University of Michigan for guidance and expertise in the area of pituitary gland development. We would like to thank the National Institute of Diabetes and Digestive and Kidney Diseases and the National Heart, Lung, and Blood Institute for financial support of these studies.

