## Supplementary figures and images for "A *Bbs5* mouse model reveals pituitary cilia contributions to developmental abnormalities"

### Supplemental Figure 1.

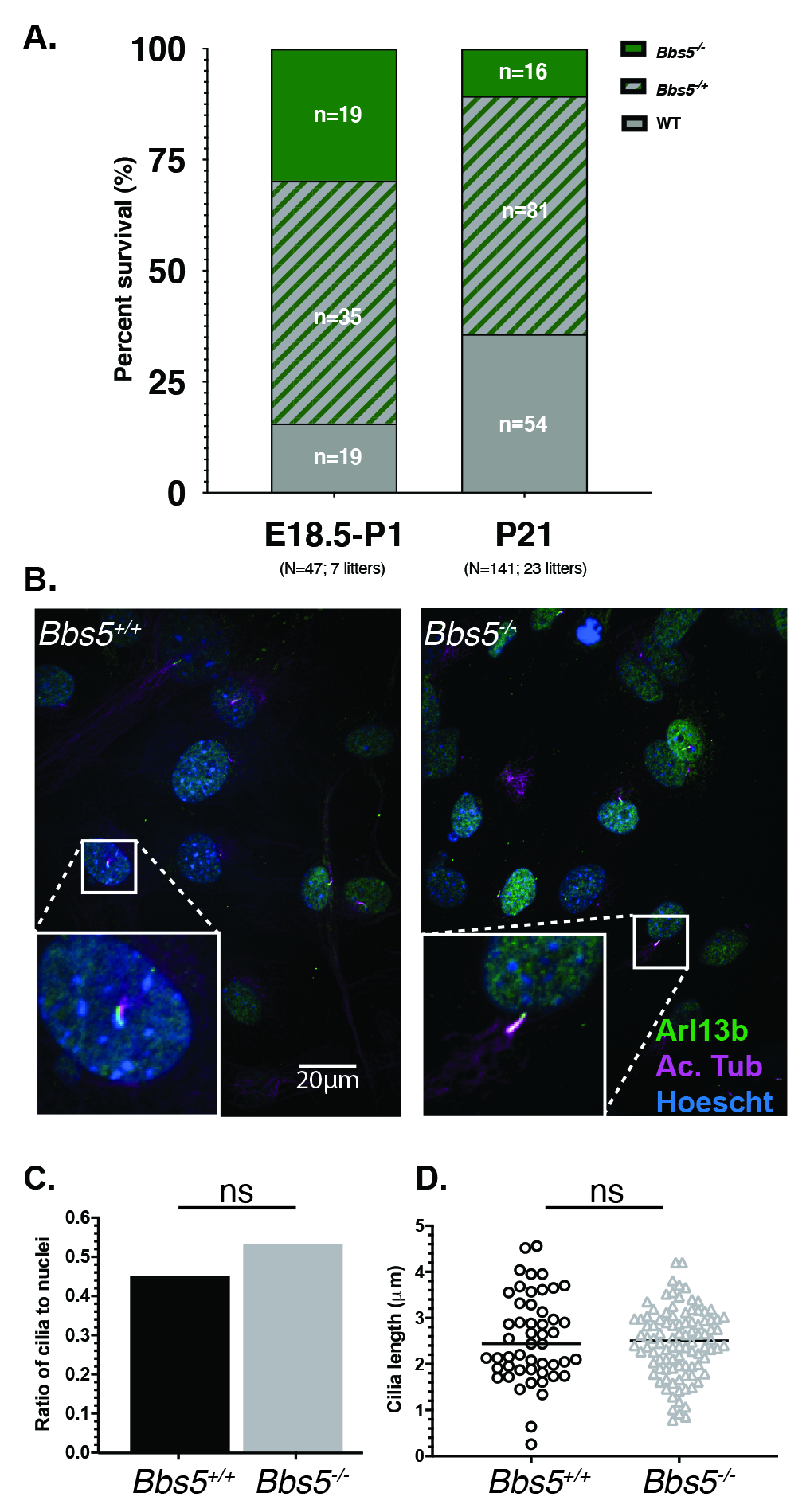

### Supplemental Figure 2.

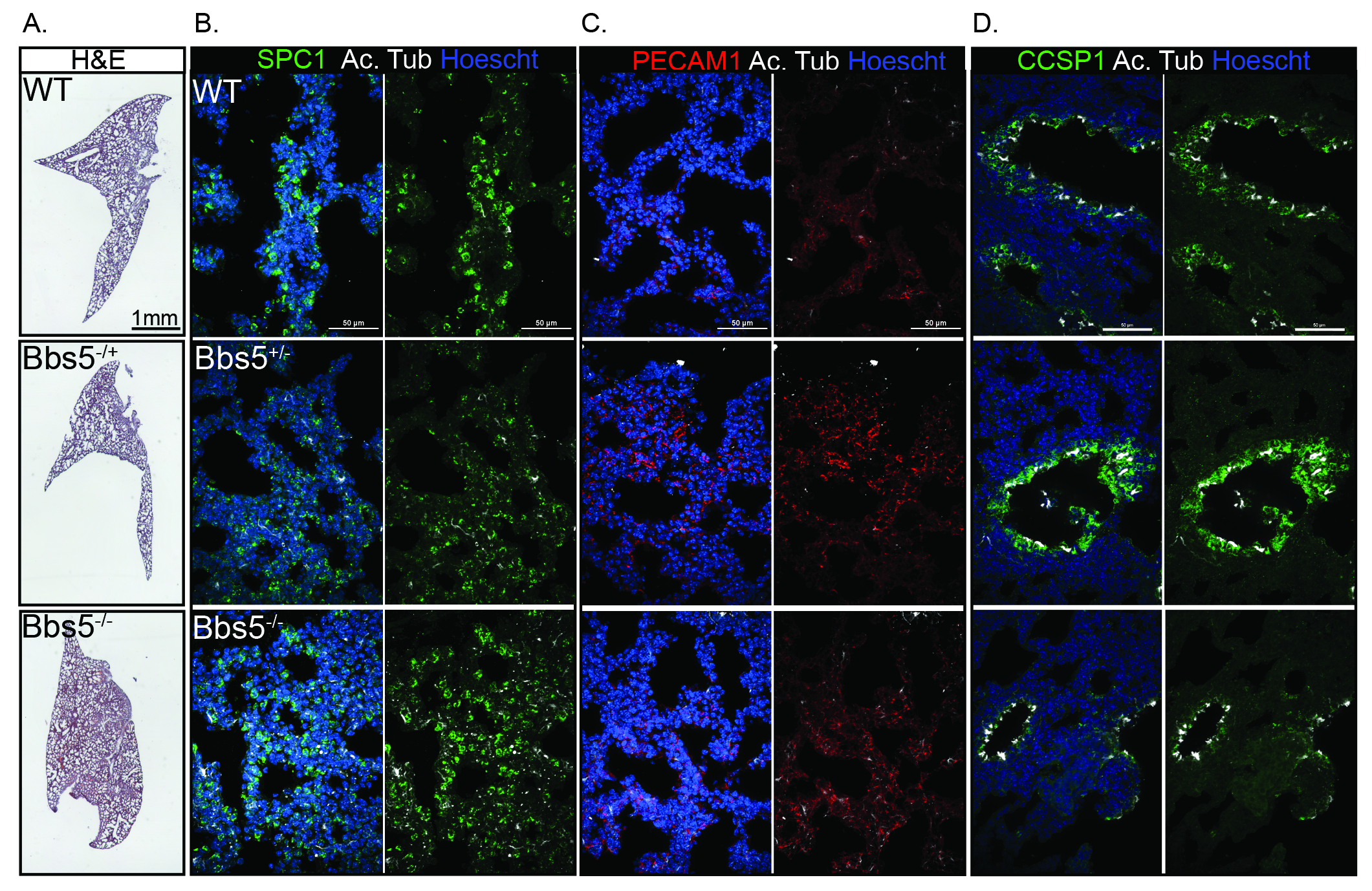

### Supplemental Video 1.

#### Slide 1
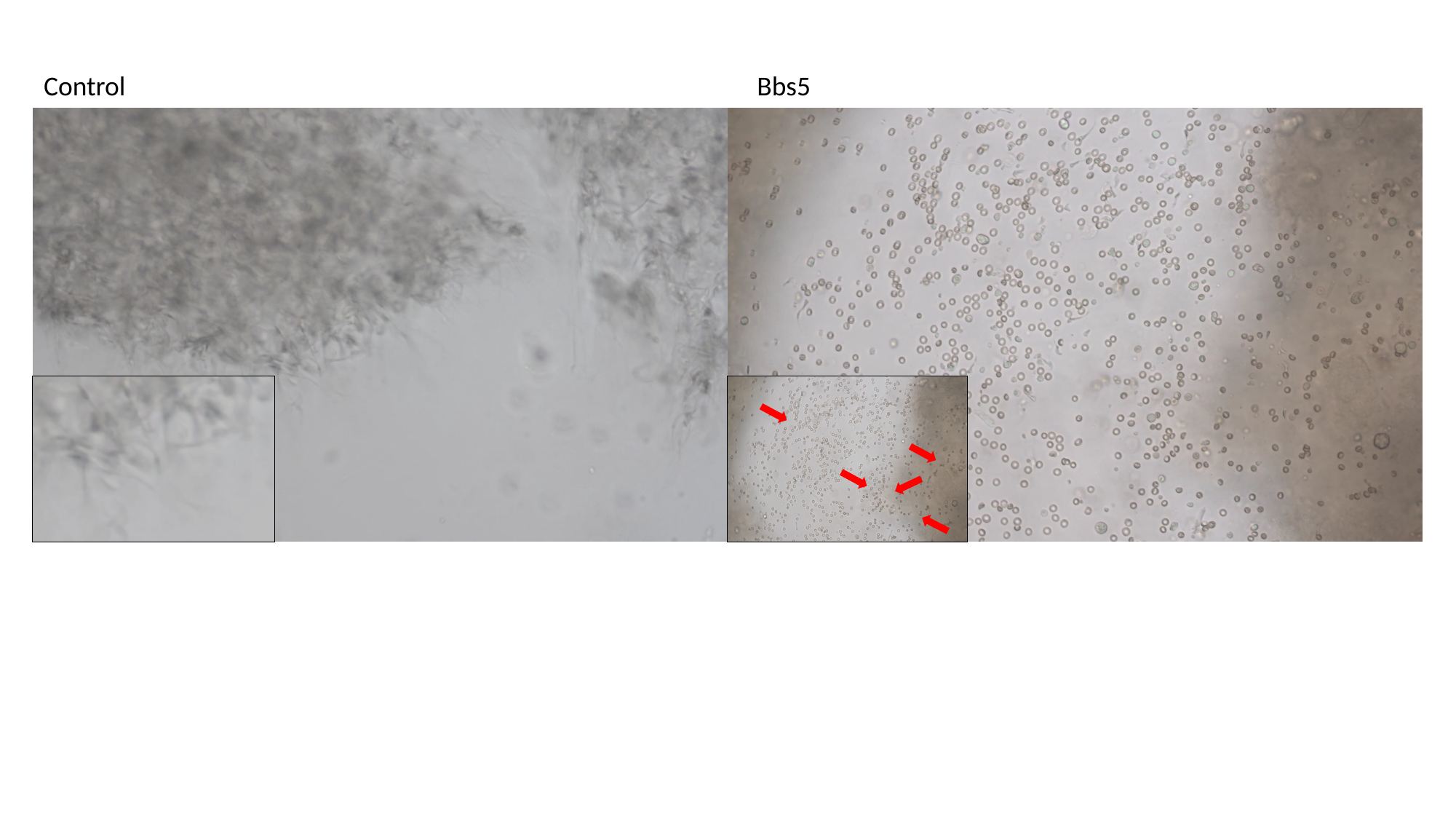

Control

### Supplemental Video 2

#### Slide 1
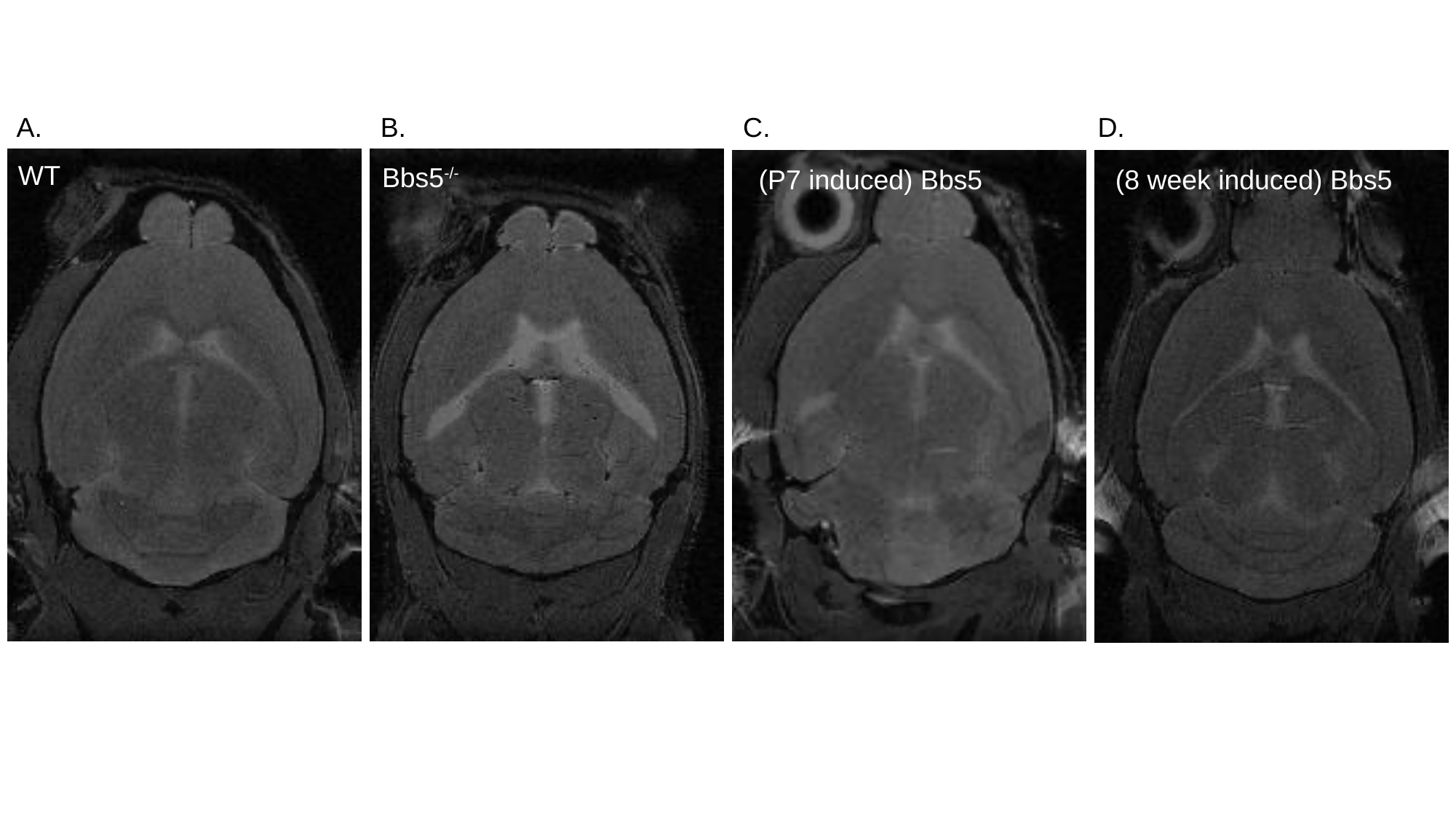

A.
B.
C.
D.
WT
Bbs5^-/-^
